# High-fidelity annotated genome of the polyploid and quarantine root-knot nematode, *Meloidogyne enterolobii*

**DOI:** 10.1101/2024.03.01.582926

**Authors:** Marine Poullet, Hemanth Gopal, Corinne Rancurel, Marine Sallaberry, Celine Lopez-Roques, Joanna Lledo, Sebastian Kiewnick, Etienne GJ Danchin

## Abstract

Root-knot nematodes of the genus *Meloidogyne* are obligatory plant endoparasites that cause substantial economic losses to the agricultural production and impact the global food supply. These plant parasitic nematodes belong to the most widespread and devastating genus worldwide, yet few measures of control are available. The most efficient way to control root-knot nematodes (RKN) is deployment of resistance genes in plants. However, current resistance genes that control other *Meloidogyne* species are mostly inefficient on *M. enterolobii*. Consequently, *M. enterolobii* was listed as a European Union quarantine pest implementing regulation. To gain insight into the molecular characteristics underlying its parasitic success, exploring the genome of *M. enterolobii* is essential. Here, we report a high-quality genome assembly of *Meloidogyne enterolobii* using the high-fidelity long-read sequencing technology developed by Pacific Biosciences, combined with a gap-aware sequence transformer, DeepConsensus. The resulting genome assembly spans 273 Mbp with 556 contigs, a GC% of 30 ± 0.042 and an N50 value of 2.11Mb, constituting a useful platform for comparative, population and functional genomics.

## Background & Summary

Root-knot nematodes (RKN) belong to the genus *Meloidogyne*, and are among the most destructive plant-parasitic nematodes^1^. Due to their extensive geographic distribution and ability to infest a wide range of host plants, they have a detrimental impact on the yield and quality of numerous economically valuable crops^2^. At present, the Meloidogyne genus comprises more than 100 described species. However, *M. arenaria, M. incognita, M. javanica and M. hapla* are considered the most widespread and damaging, RKN species^3^. In recent years, *M. enterolobii* has received increasing attention due to its unique ability to overcome several sources of resistance against RKN^2,4,5^.

The species *Meloidogyne enterolobii* was originally first described as *M. incognita* from a population obtained from the Pacara Earpod Tree (*Enterolobium contortisiliquum* [Vell.] Morong) in Hainan Island, China by Yang and Eisenback (1983)^6^. Later in 1988, Rammah and Hirschmann described a new species^7^, *M. mayaguensis*, sampled from eggplant (*Solanum melongena* L.) roots from Puerto Rico. However, this species was later synonymized with *M. enterolobii*, based on the same esterase phenotype and mitochondrial DNA sequence^8,9^.

*Meloidogyne enterolobii* has an extremely high damage potential^10^, surpassing many of the other root-knot nematode species studied so far^11,12^. The reports of severe damage in high-value crops have increased in the past years^13,14^. In 2009, the European Plant Protection Organization (EPPO) performed a risk analysis, which came to the conclusion that this species was recommended for regulation and placed on the EPPO A2 list in 2010. Following numerous interceptions over the years, it was concluded that *M. enterolobii* fulfilled the conditions provided in Article 3 and Section 1 of Annex I to Regulation (EU) 2016/2031 in respect of the Union territory and therefore should be listed in Part A of Annex II to Implementing Regulation (EU) 2019/2072 as Union quarantine pest^15^. However, once damage is detected, *M. enterolobii* identification is challenging due to morphological resemblances it shares with other root-knot nematode species^11,14,16,17^.

In that perspective, providing high-quality nuclear and mitochondrial genomes for this species can accelerate the development of reliable molecular markers and the understanding of the biology of *M. enterolobii*. A first version of the genome of *M. enterolobii* was published in 2017 as part of a comparative genomics analysis with other root-knot nematodes^18^. The population named L30, originated from Burkina Faso and was sequenced using Illumina short reads. Consequently, the assembled genome was quite fragmented with >46,000 contigs and an N50 length <9.3kb, precluding analyses of structural variants or conserved synteny with other *Meloidogyne* species. Nevertheless, this initial genome allowed confirming that this species was likely polyploid, similarly to other tropical parthenogenetic root-knot nematodes^19^. More recently, a *M. enterolobii* genome assembled from PacBio RS long reads and polished with Illumina short reads was published^20^. The sequenced population named Mma-II, was isolated from infected tomatoes in a Swiss organic farm^21^. With <4,500 contigs and an N50 length of 143kb, the genome assembly represented a substantial improvement compared to the only other assembly available at this time. However, recent progress in the quality and data volume of long-read sequencing technologies^22^ promises even more contiguous and higher-quality genomes even for complex polyploid species and including the Meloidogyne genus^23,24^. Therefore, we used the PacBio HiFi, highly accurate long-read sequencing technology to produce a more contiguous and reliable reference genome for this quarantine plant-parasitic nematode.

Using this technology and further improvement of the quality of the reads, we assembled the genome of the *Meloidogyne enterolobii* population (E1834), originally isolated from the roots of eggplant collected in Puerto Rico, in 556 contigs with an N50 length surpassing 2Mb and a genome assembly size of 273Mb, consistent with previous flow cytometry estimation on a population from Guadeloupe island (274.7 +- 18.52 Mb)^20^. Compared to the previous long-read version of the genome, this constitutes an improvement of the N50 contiguity by more than one order of magnitude, with the number of contigs divided by almost 10.

Further quality check of our genome assembly and comparison with previous assemblies confirmed the correct species identification for population E1834 and for the isolate from Burkina Faso previously sequenced with short reads. However, our study also revealed that the Mma-II Swiss population previously sequenced with PacBio RS underwent a contamination by *M. incognita,* which over several generations in a greenhouse, completely overtook originally described *M. enterolobii* population M-ma-II. As this population was not maintained as a single egg mass line, contamination by a highly virulent and equally pathogenic *M. incognita* population remained undetected. Mis-identification among *Meloidogyne* species is not uncommon as reported populations of *M. ethiopica* in Europe were later identified as *M. luci*^25^. Consequently, the genome assembly in that publication^20^ corresponded to *M. incognita* implying no long-read-based contiguous genome for *M. enterolobii* was finally available so far. This finding also motivated us to develop a methodology based on mitochondrial genomes reconstruction and relative coverage to detect contamination between closely related species which are not detectable with standard Blobtools^26^ approaches based on contigs GC content and coverage. This methodology can be reused to confirm correct species identification in sequencing projects.

Overall, we propose a high-quality contiguous genome for *M. enterolobii* constituting a reliable resource for within- and between-species comparative genomics. The contiguity of the genome enables study of structural variations and conserved synteny, which will be essential towards comprehensive identification of genomic variations in relation with the host range of this quarantine nematode species in Europe.

## Methods

### Nematode collection and DNA extraction

The *Meloidogyne enterolobii* population (E1834) was originally isolated from the roots of eggplant collected in Puerto Rico and has been maintained since 2005 in the *Meloidogyne* sp. reference collection at The Netherlands Institute for Vectors, Invasive plants and Plant health (NIVIP) Wageningen, Netherlands. In 2020, this population was kindly provided by NRC, for the research conducted in the framework of the project AEGONE (No. 431627824r) and has been maintained at the Julius Kühn Institut (JKI) in Braunschweig, Germany in a greenhouse on the resistant tomato cultivar ‘Phantasia’. Nematodes used for DNA extraction were obtained from single egg mass (SEM) lines. To obtain these lines, 12 single females with egg masses were carefully picked from the infected roots of tomato and second stage juveniles (J2) were allowed to hatch in six well plates (SARSTEDT AG & Co. KG, Nümbrecht, DE) with 5ml molecular grade water per well. After one week at room temperature (20 ± 1°C) in the dark, 10 wells with the highest number of hatched J2s were selected for inoculation. In addition, two J2s from each egg mass were collected for DNA extraction and species verification by Real-time PCR^27^ and SCAR species-specific markers^28,29^. For multiplication of the SEMLs, five-week-old tomato seedlings from the cultivar ‘Phantasia’ were transplanted into 1000 ml clay pots (Risa Pflanzgefässe GmbH, Germany) containing 750ml quartz sand (0.3-1mm) supplemented with slow-releasing fertilizer, Osmocote (1.5g/L). Afterwards, tomatoes were inoculated with J2s obtained from the respective egg masses. Tomato plants were maintained in a greenhouse at 20 to 25°C with 16h of light and 8h of darkness. Plants were watered daily and fertilized once per week with Wuxal® super solution (8:8:6; N: P: K, Hauert MANNA, Nürnberg, DE). After 8 weeks, the galled roots were carefully washed free of sand and the eggs and juveniles (E&J) were extracted with 0.7% chlorine solution^30^. The resulting E&J suspension was counted to identify the line with the highest reproduction rate. The SEML number 4 was therefore selected (Table 1) for further experiments and production of DNA.

**Table 1:** Number of newly produced eggs and juveniles per root system of 10 single eggs mass lines of *Meloidogyne enterolobii* population (E1834). Tabulated values are the mean count of Eggs and Juveniles with standard error for different SEML.

|  | Number of Egg & Juveniles<br>per root |
| --- | --- |
| SEML 1 | 43,200 ± 577.35 |
| SEML 2 | 22,7800 ± 1285.82 |
| SEML 3 | 56,600 ± 1604.16 |
| SEML 4 | 454,800 ± 1442.22 |
| SEML 5 | 230,400 ± 945.16 |
| SEML 6 | 3000 ± 503.32 |
| SEML 7 | 187,800 ± 2457.64 |
| SEML 8 | 170,000 ± 416.33 |
| SEML 9 | 109,000 ± 1222.02 |
| SEML 10 | 189,800 ± 901.85 |

### DNA extraction

The selected SEML 4 was multiplied on tomato plants to obtain J2 for DNA extraction. Galled tomato roots were carefully washed free of sand and placed in a mist chamber to collect freshly hatched J2 after 14 days. The J2 suspension was purified by the modified centrifuge-floatation method^31^ with a 45% sugar solution to reduce contaminations such as root debris, bacteria, fungal spores, etc… Afterwards, approximately 50,000-70,000 J2 were transferred into 1.5ml Eppendorf tube and washed 3 times with molecular grade water. After freezing in liquid nitrogen, DNA was extracted from the homogenized sample using the MasterPure Complete DNA & RNA Purification Kit (Lucigen) following the manufacturer’s protocol. The DNA was suspended in 10mM Tris-HCl buffer and the DNA concentration was determined with either a Qubit^TM^ 4 fluorometer (Life Technologies, Singapore) or NanoDrop 2000 ^TM^ spectrophotometer (Thermo Fisher Scientific, USA). The NanoDrop 2000™ was blanked using the respective elution buffer for the method. DNA concentration was measured using Qubit™ (1X dsDNA HS (High Sensitivity) Assay Kit, Invitrogen, #Q32853) and NanoDrop 2000™. Purity was measured using the 260/280 nm and 260/230 nm absorbance ratios of NanoDrop 2000™.

### Genome sequencing and read processing

The long-fragment DNA libraries from the *M*. *enterolobii* population E1834 were constructed at the GeT-PlaGe core facility, INRAE Toulouse according to the manufacturer’s instructions “Preparing whole genome and metagenome libraries using SMRTbell® prep kit 3.0”. At each step, DNA was quantified using the Qubit dsDNA HS Assay Kit (Life Technologies). DNA purity was tested using the nanodrop (Thermofisher) and size distribution and degradation assessed using the Femto pulse Genomic DNA 165 kb Kit (Agilent). Purification steps were performed using AMPure PB beads (PacBio) and SMRTbell cleanup beads (PacBio). A DNA damage repair step was performed using the « SMRTbell Damage Repair Kit SPV3 » (PacBio). A total of 9.4μg of DNA was purified and then sheared at 20 kb using the Megaruptor system (Diagenode). Using SMRTbell® prep kit 3.0, a Single strand overhangs removal, a DNA and END damage repair step were performed on 8.3μg of sample. Subsequently, blunt hairpin adapters were ligated to the library and a nuclease treatment was performed using the nuclease mix of “SMRTbell® prep kit 3.0”. A size selection step using a 6kb cutoff was performed on the BluePippin Size Selection system (Sage Science) with “0.75% DF Marker S1 6-10 kb vs3 Improved Recovery” protocol. Using Binding kit 2.2 kit and sequencing kit 2.0, the primer V5 annealed and polymerase 2.2 bounded library was sequenced by diffusion loading with the adaptive loading method onto 1 on Sequel II instrument at 90pM with a 2 hours pre-extension and a 30 hours movie.

The Sequel II sequencing system outputs 1To of raw data into a subread file. This contains unaligned base calls from high-quality regions, the complete set of base quality values and kinetic measurements from the sequencing instrument. This subread file is used as input for the Circular Consensus Sequencing (CCS v6.4.0) analysis to generate a draft consensus sequence. Very low-quality reads (<Q9) were filtered out by using the parameter --min-rq=0.88. To further improve the quality of the PacBio Sequel II reads, we have used a gap-aware sequence transformer, DeepConsensus^32^ (v1.1.0). As a final step, the previous subreads were aligned to the draft consensus sequence using ACTC^33^ with default parameters (v0.2.0) and used as input to the DeepConsensus transformer-encoder. The Phred-scale read accuracy score (Qconcordance) has been calculated according to Baid et al.^32^ where Qconcordance = - 10*log10 (1-identity) and identity = matches / (matches + mismatches + deletions + insertions).

### Ploidy, heterozygosity, and genome size estimation

To infer the ploidy level of the *M. enterolobii* population E1834, a k-mer-based approach was employed to profile the genome. The k-mer frequencies in DeepConsensus sequencing reads were analyzed using KMC^34^ (v3.0.0, kmc -k21 -m100 -ci1 -cs10000). In accordance with the author’s recommendations, canonical 21-mers were extracted using a hash and organized in a histogram file using the kmc_tools transform option. To determine the appropriate coverage thresholds required for the inference, the KMC histogram file is utilized as input for the cutoff option in Smudgeplot^35^ (v0.2.4). Subsequently, we generated a smudge plot using the coverage of the identified k-mer pairs to determine ploidy.

To estimate genome size and heterozygosity prior to assembly, we used Genomescope^35^ (v2.0) on the histogram file generated from Jellyfish^36^ (v2.3.0, jellyfish histo -h 1000000) and the coverage thresholds produced by the Smudgeplot cutoff tool. The final genome size is therefore obtained by multiplying the haploid genome size by the previously estimated ploidy level.

### Genome assembly, estimation of completeness and contamination

DeepConsensus reads were trimmed of remaining adapters using HifiAdapterFilt^37^ (v2.0.0) with default parameters. The trimmed reads were then used as input to the Peregrine-2021 assembler^38,39^ (v0.4.11) while increasing the default number of best overlaps for each initial graph (parameter --bestn 8). This parameter is optimized for highly heterozygous genomes. To further assess genome assembly completeness in a reference-free approach, we used Merqury’s algorithm^40^ (v1.3). This tool uses k-mer frequencies to evaluate a genome’s base accuracy and completeness. This is achieved by counting and comparing the distribution of canonical 21-mers found in the assembled genome with those detected in the high-accuracy DeepConsensus read set. Merqury’s k-mer analysis will therefore indicate whether the genome assembly has captured all the information present in the HiFi reads.

The screening of the contig assembly for potential contaminants by non-nematode sequences was done with the Blobtools^26^ pipeline (v3.2.6). DeepConsensus polished long-reads were aligned to the contigs with Minimap2^41^ (v2.24) and the map-hifi parameter. Each contig was then assigned to a taxonomic group based on the BLAST^42^ (v2.13.0+) analysis results against the NCBI nucleotide (nt) database^43^. Particular attention was paid to contigs of non-nematode taxa or contigs with a GC percentage deviating from the average GC content (around 30%^18^) of the *M. enterolobii* population E1834 to detect possible contamination. A total of 39 contigs spanning ∼2.5 Mbp were discarded from the assembly. The resulting assembly of 556 contigs is used for downstream analyses.

### Mitochondrion assembly and functional annotation

The circular mitochondrial genome sequence was reconstituted using the ALADIN^44^ package (v1.1) and DeepConsensus HiFi reads in input with default parameters. We employed as a reference seed sequence the complete mitochondrion of *M. enterolobii* previously downloaded from the GenBank database (BioProject: PRJNA927338^45^). The annotation was carried out using GeSeq^46^, encompassing both the tRNA, the rRNA and the protein-coding genes. We set the minimum threshold of 85% for the protein and non-coding DNA search identity, and we used seven Meloidogyne mitochondrial genomes as third-party references (Table 2). The tRNAs prediction was also performed using third-party predictors, such as tRNAscan-SE^47^ (v2.0.7), ARAGORN^48^ (v1.2.38), and ARWEN^49^ (v1.2.3), with codon usage corresponding to Metazoan and Invertebrate Mitochondrial.

**Table 2:** Mitochondrial sequence data statistics for different nematodes and their correspondence with the modified Blobtools analyses. Seven Meloidogyne mitochondrial sequences have been analyzed for this study. For each species, a specific custom TaxID corresponding to a different phylum has been used.

| Species | Length (bp) | GenBank accession | TaxIDs | Custom TaxIDs |
| --- | --- | --- | --- | --- |
| *M. graminicola*<sup>55</sup> | 19589 | NC_056772 | 189291 | 4890(Ascomycota) |
| *M. arenaria*<sup>56</sup> | 17580 | NC_026554 | 6304 | 6340 (Annelida) |
| *M. enterolobii*<sup>45</sup> | 17053 | NC_026555 | 390850 | 390850 |
| *M. javanica*<sup>57</sup> | 18291 | NC_026556 | 6303 | 10190 (Rotifera) |
| *M. incognita*<sup>58</sup> | 17662 | NC_02409 | 6306 | 6656 (Arthropoda) |
| *M. chitwoodi*<sup>58</sup> | 18201 | KJ476150 | 59747 | 6447 (Mollusca) |
| *M. oryzae*<sup>59</sup> | 17066 | MK507908 | 325757 | 7711 (Chordata) |

### Gene prediction and genome structure determination

Gene models prediction was done with the fully automated pipeline EuGene-EP^50^ (v1.6.5). EuGene has been configured to integrate similarities with known proteins of *Caenorhabditis elegans* (PRJNA13758) from WormBase Parasite^51^ and “nematoda” section of UniProtKB/Swiss-Prot library^52^, with the prior exclusion of proteins that were similar to those present in RepBase^53^. The dataset of *Meloidogyne enterolobii* transcribed sequences^20^ was aligned on the genome and used by EuGene as transcription evidence. Only the alignments of datasets on the genome spanning 30% of the transcript length with at least 97% identity were retained. The EuGene default configuration was edited to set the “preserve” parameter to 1 for all datasets, the “gmap_intron_filter” parameter to 1 and the minimum intron length to 35 bp. Finally, the Nematodes-specific Weight Array Method matrices were used to score the splice sites (available at this URL: http://eugene.toulouse.inra.fr/Downloads/WAM_ nematodes_20171017.tar.gz).

Genome structure analysis was conducted using MCScanX^54^, with default settings. First, the whole proteome of the *M. enterolobii* population E1834, predicted by EuGene, was self-blasted with an E-value cutoff of 1e-25, a maximum of 5 aligned sequences, and maximum 1 high-scoring pair (hsp). Subsequently, we used gene location information extracted from the GFF3 annotation file of EuGene, along with homology information based on the all-versus-all BLASTP analysis, to identify and categorize each duplicated protein-coding gene into one of five groups using the duplicate_gene_classifier program implemented in the MCScanX package. These groups are: singleton, proximal, tandem, whole-genome or segmental duplications (WGD), and dispersed duplications. Singleton refers to cases where no duplicates are found in the assembly. Proximal duplicates refer to gene duplications that are on the same contig and separated by 1 to 10 genes. Tandem duplicates, on the other hand, are consecutive. WGD are identified when they form collinear blocks with other pairs of duplicated genes. Finally, dispersed duplicates are those that cannot be assigned to any of the above-mentioned categories.

### Species verification and validation

In the following step, we further screened the genomic reads for potential contamination, this time by other root-knot nematode sequences. Blobtools allows the identification of potential contamination in genome assemblies, but only at distant taxonomic levels between different phyla (e.g., Chordata, Nematoda, Arthropoda, …). Therefore, although contamination can be detected and cleaned at this level, it remains undetectable at the intra-phylum level (e.g. within Nematoda). To allow the detection of contamination by other closely related nematodes at the reads level, we adapted the Blobtools pipeline to work with mitochondrial genomes. The polished long-reads were aligned against complete mitochondrial sequences for seven *Meloidogyne* species downloaded from the NCBI database (Table 2), using the same procedure as above. Since Blobtools only works at the phylum and not species rank, we used a script to create an additional hits file and assign a custom NCBI phylum TaxID to each species. The seven *Meloidogyne* samples have been then temporarily assigned to a different phylum for the BlobPlot visualization only (Table 2).

Species-specific SCAR (sequence characterized amplified region) markers are routinely used to confirm species identity in plant-parasitic nematodes^60^. SCAR markers are locus-specific fragments of DNA that are amplified by PCR using specific 15-30bp primers. In this study, we retrieved primer sequences of species-specific SCAR markers from the literature (Supplementary Table 1) for four Meloidogyne species with genome assemblies publicly available and belonging to the same clade (*M. arenaria*, *M. incognita*, *M. javanica* and *M. enterolobii*). We aligned all the primers to all the above-mentioned genomes with BLAST and when the primer pairs matched on the same contig, we retrieved from the genome the ‘virtual’ PCR products. After verification of consistency with the lengths from the literature, the virtual PCR products were then aligned to the two previous and present versions of *M. enterolobii* genome assemblies with an E-value threshold of 1e-25.

## Data Records

All the PacBio HiFi sequence data as well as the genome assemblies and gene predictions supporting the results of this paper have been deposited and are publicly available at the EMBL-EBI’s European Nucleotide Archive (ENA) under accession number PRJEB69523 (https://www.ebi.ac.uk/ena/browser/view/PRJEB69523)^61^. All the processed data, including genome assemblies^62^, gene predictions^63^, and all the structural annotation^64^ results have been deposited and are publicly available at the Recherche Data Gouv institutional collection.

## Technical Validation

### Assessing read accuracy

After implementing the DeepConsensus^32^ sequence transformer procedure, statistical analysis showed an increase in the number of high-quality reads obtained (Fig. 1, Table 3) with long fragment DNA reads of up to 26kb in length. The average length of the reads is around 11kb with a total number of 2.4M reads, and a higher average Phred-scale read accuracy score (Qconcordance), which increased from 31.95 before to 34 after DeepConsensus. This transformer has elevated the PacBio HiFi read yield to a minimum Q30 by 10% and a minimum Q40 score (99.99% read accuracy) by 70%. Furthermore, we have retrieved 198,880 long-reads that were initially dismissed prior to treatment in the filter, providing us with more chances to comprehensively capture the entire genome of *M. enterolobii*.

**Figure 1.**
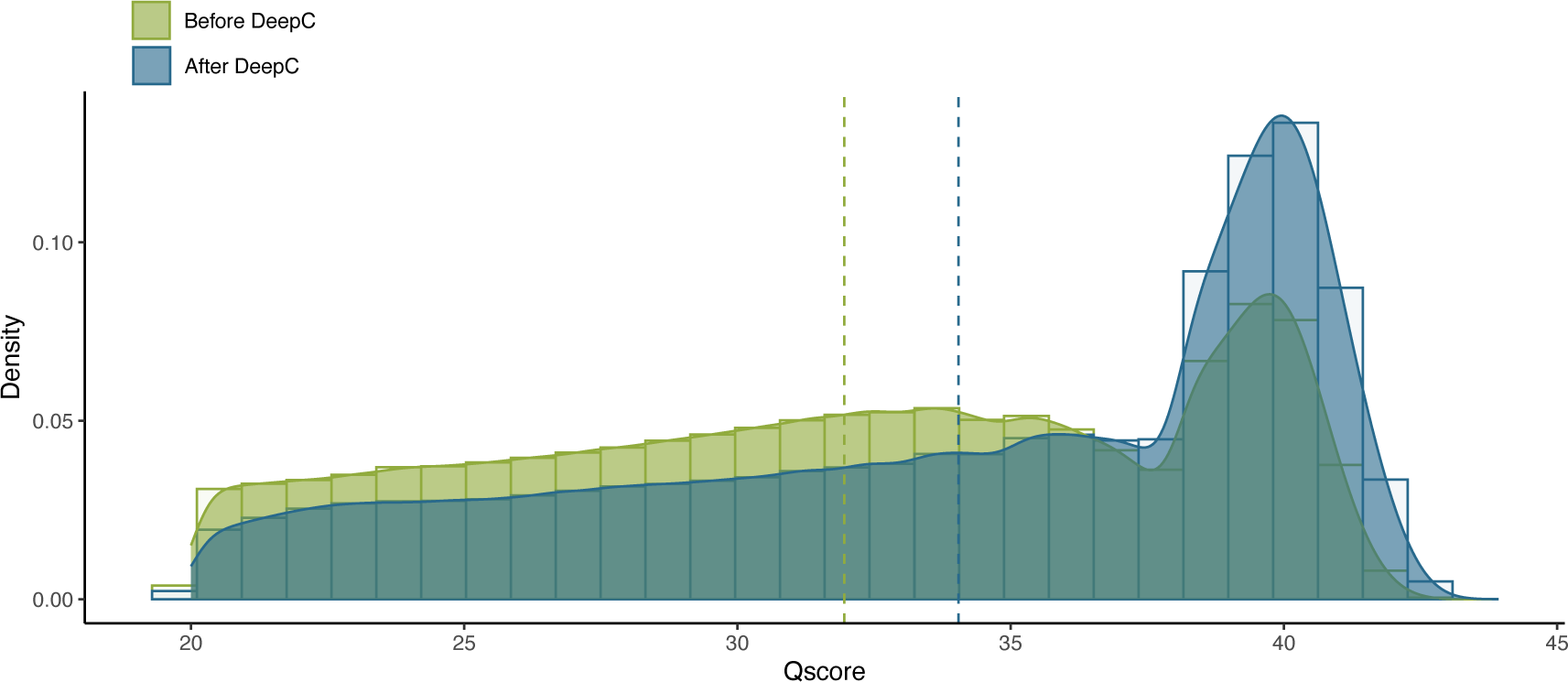
Distribution of raw PacBio HiFi reads before and after DeepConsensus treatment. Comparison of the Concordance Qscore before and after DeepConsensus. The average phred-scale read accuracy score has increased by two points after treatment.

**Table 3:** Read statistics before and after the use of DeepConsensus sequence transformer. After DeepConsensus treatment, a higher number of reads with higher quality have been retrieved.

|  | Before DeepConsensus | After DeepConsensus |
| --- | --- | --- |
| Longest read | 26 302 bp | 26 302 bp |
| Mean length | 11 194 bp | 11 187 bp |
| Number of reads | 2 250 199 | 2 449 079 |
| Number of bases | 25 442 730 700 | 27 277 356 063 |
| Average Qscore | 31.95 | 34.04 |

### Profiling genome ploidy level, heterozygosity, and size

Prior to assembling a genome, it is crucial to evaluate its ploidy and size. The distribution of k-mer frequencies within the DeepConsensus sequencing reads allows estimating major genome features such as ploidy level, genome size, and heterozygosity rate. As GenomeScope2^35^ can only precisely examine organisms when a definite ploidy is known, we utilized first, the results of Smudgeplot^35^ (Fig. 2a) to provide GenomeScope2 with estimated ploidy level. Each smudge on the graph appears to be distinct, indicating sufficient sequencing coverage for further analysis. The most prevalent smudge corresponds to a predicted triploid AAB genome for the *M. enterolobii* population E1834 (Table 4). This result is consistent with previous k-mer analysis performed on the short-reads for the L30 population from Burkina Faso^18,65^.

**Figure 2.**
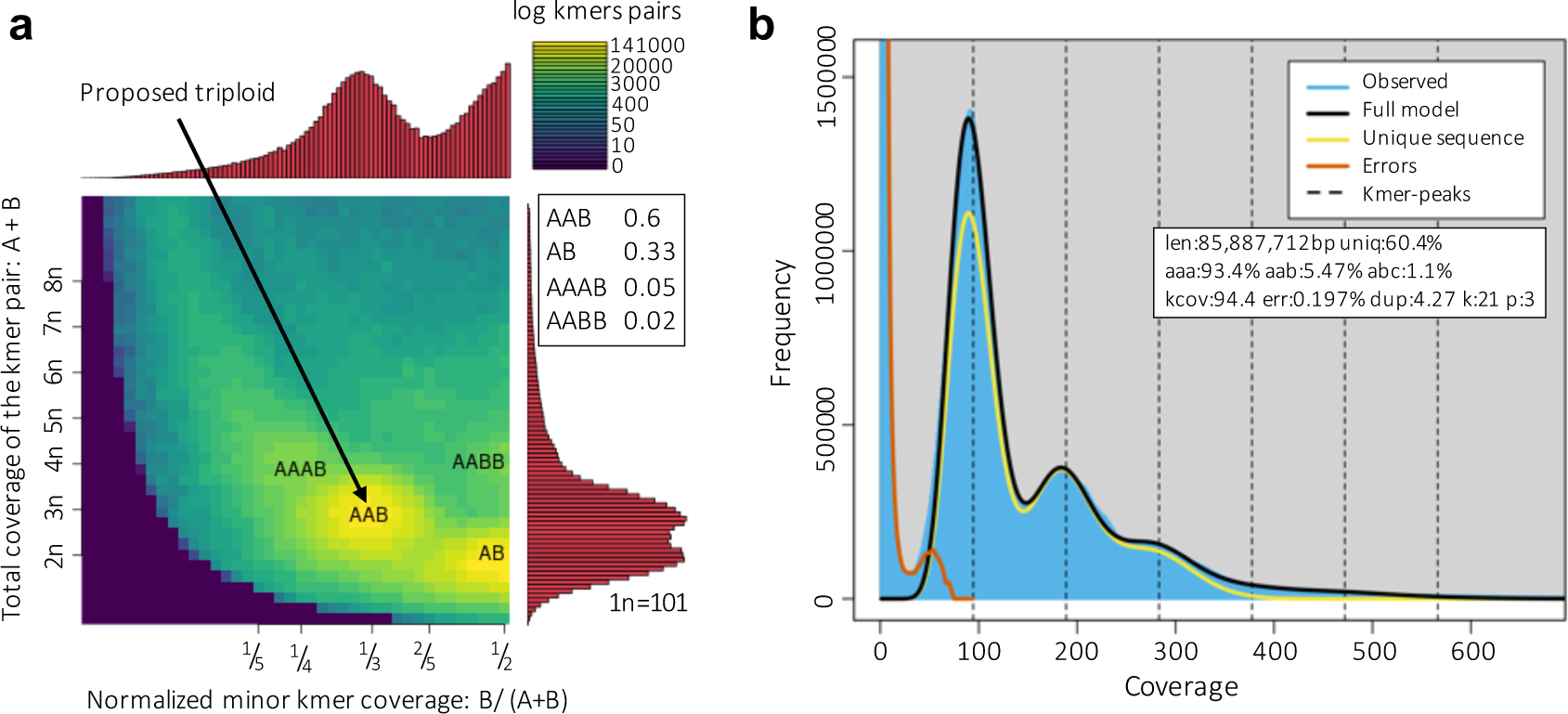
Genome profiling of *M. enterolobii*. **A**. Smudgeplot of *M. enterolobii* extracting 21-mers from DeepConsensus reads. The color intensity of each smudge reflects the approximate number of k-mers per bin. *M. enterolobii* E1834 population is proposed as a triploid organism. **B**. GenomeScope2 k-mer profile and estimated parameters for the triploid nematode *M. enterolobii*. Coverage (kcov), error rate (err.), haploid genome size estimation (len.), k-mer size (k) and ploidy level (p). The peak heights are proportional to the species’ heterozygosity. *M. enterolobii* shows a high heterozygosity.

**Table 4.** Summary of peaks detected by Smudgeplot. This result proposes that the *M. enterolobii* E1834 population is triploid.

| Peak | Kmers # | Kmers proportion | Summit B/ (A+B) | Summit A+B |
| --- | --- | --- | --- | --- |
| AAB | 5 952 405 | 0.60 | 0.34 | 309.70 |
| AB | 3 243 405 | 0.33 | 0.49 | 191.27 |
| AAAB | 508 560 | 0.05 | 0.24 | 451.82 |
| AABB | 213 490 | 0.02 | 0.49 | 404.45 |

Subsequently, we estimated the genome size using GenomeScope2 with a ploidy level of 3 (Fig. 2b). The genome size was determined by multiplying the estimated haploid genome length (85,887,712 bp) by the previously estimated ploidy level (p=3), providing an estimated genome size of 257.66 Mb.

Furthermore, the GenomeScope2 k-mer histogram of this polyploid population displays a distinct multimodal profile, with a substantial first peak located at roughly 95X, a smaller second peak at about 187X, and finally, an additional peak at 282X, typical for triploid genomes. Finally, GenomeScope2 estimated a highly heterozygous genome (6.5 % estimated on average), consistent with a previous estimation of ca. 6.1% on the L30 population from Burkina Faso^18^. It should be noted that the term heterozygosity does not exactly apply here as we do not measure divergence between homologous chromosomes in a diploid genome but between the AAB subgenomes in a triploid species. Therefore, we will refer to average nucleotide divergence between subgenomes in the rest of the manuscript.

### *De novo* genome assembly

After filtering and elimination of the contaminated and mitochondrial contigs, the resulting genome of the *Meloidogyne enterolobii* population E1834 is assembled in 556 contigs with a total size of 273 Mbp. The corresponding contig N50 length is equal to 2.11Mb, with the longest being 8.3 Mb, long.

The genome assembly size is congruent with the total DNA content estimated by flow cytometry (274.69 ± 18.52 Mb) on another population of *M. enterolobii* from Guadeloupe island^20^, indicating that the assembly represents a complete *M. enterolobii* genome.

However, the genome size estimated by analysis of k-mer distribution (257.66 Mb) is lower than the assembly size and in the lower range of the flow cytometry evaluation. A previous study showed that the accuracy of genome size estimation based on k-mer frequencies can be affected by repeats, high heterozygosity and sequencing errors^66^. This suggests that the high heterozygosity rate or repeat-richness in the *M. enterolobii* genome could have played a role in this underestimation.

To further assess genome assembly quality metrics and evaluate genome’s base accuracy and completeness we used Merqury^40^. In the Merqury spectrum produced (Fig. 3a), the first and prominent 1-copy peak at a ∼100X multiplicity corresponds to k-mers in the reads detected only one time in the assembly. This can be interpreted as heterozygous regions between the three subgenomes. The second peak at twice this multiplicity (∼200X) corresponds to homozygous k-mers present in the reads and detected two times in the assembly. This most likely represents regions identical between two of the three AAB subgenomes. Similarly, most of the k-mers detected at 3 times in the assembly, probably represent regions identical between the three AAB subgenomes. Conversely, the grey peak at low multiplicity represents rare k-mers which solely exist within the read set and are probably due to sequencing errors. This Merqury plot indicates a lack of missing content as there is no subsequent grey peak at the 1-copy peak (∼100X coverage). Additionally, the AAB subgenomes are divergent enough to have been mostly separated (or unzipped) during the assembly, as there is no smaller second grey peak beneath the 2-copy peak at twice the coverage (∼200X). This can also be observed in the coverage plot provided by bedtools^67^ v2.29.0 (Fig. 3b), where the coverage depth for each base on each contig has been computed. We can clearly see a prominent peak located at roughly 101X corresponding to the haploid coverage found in the k-mers with Smugeplot (Fig. 2a), and a shoulder at roughly twice the haploid coverage probably represented few identical regions between sub-genomes that have not been completely unzipped during assembly.

**Figure 3.**
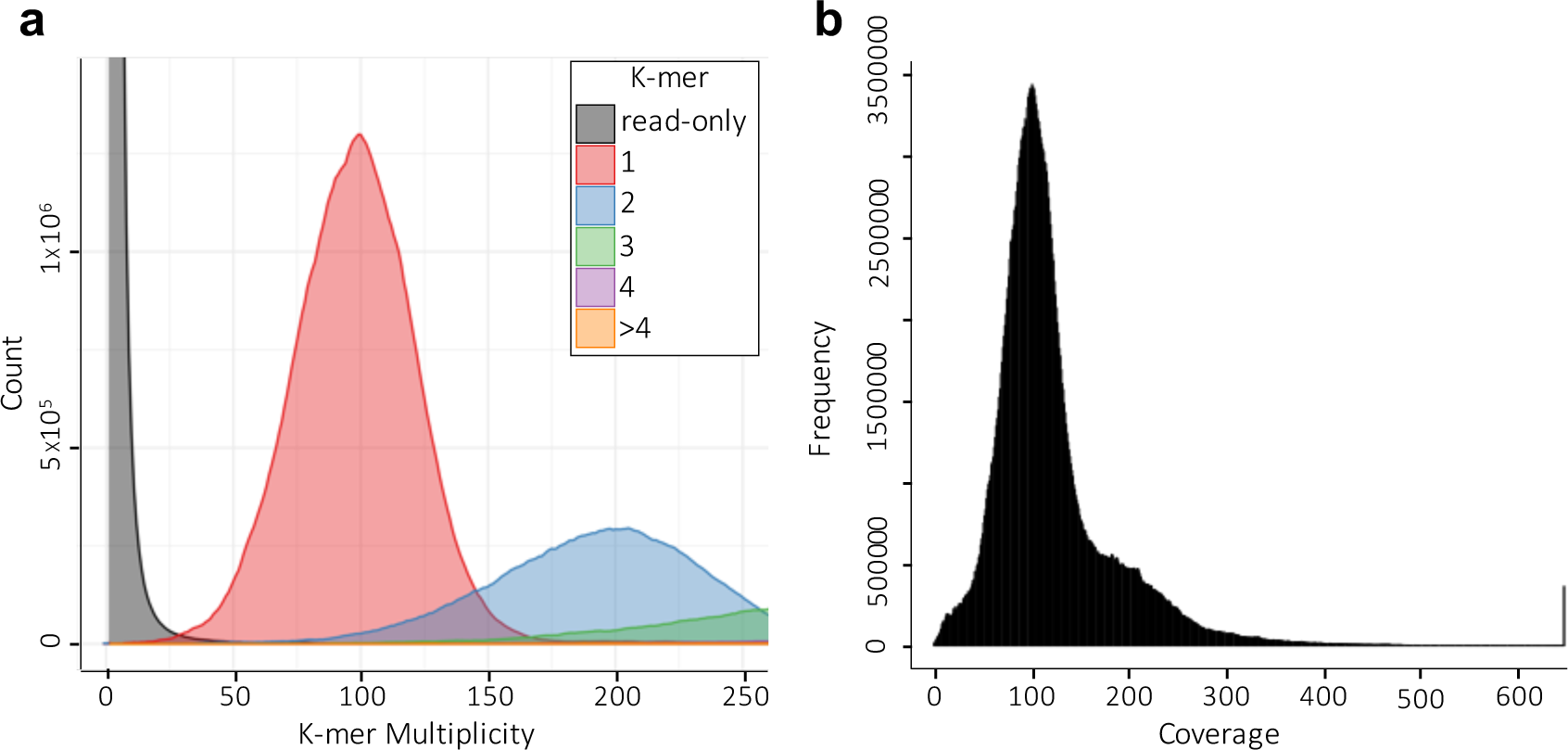
Genome assembly spectra. **A**. The Merqury spectrum plot using DeepConsensus reads tracks the multiplicity of each k-mer detected in the read set. The plot is color-coded according to the number of times a k-mer is found in an assembly. **B**. Bedtools per-base reports coverage for the assembly. The three *M. enterolobii* AAB subgenomes were effectively separated during assembly using Peregrine.

Overall, the k-mer analysis with Merqury indicates that all the information present in the HiFi reads has been captured in the genome assembly, further suggesting a complete genome.

### Validating species identity and purity

We confirmed the purity of the *M. enterolobii* population E1834 and the correct species identification by using, first, the Blobtools^26^ pipeline. The pipeline generated BlobPlots, which are two-dimensional plots depicting contigs presented as circles, whose diameters are proportional to the sequence length and are colored based on their taxonomic affiliation, determined by the BLAST similarity search results against the NCBI nt database^43^. The relative positions of the circles are according to their GC content and coverage by the long reads. Following the removal of contaminant contigs, the resulting BlobPlot is shown in Figure 4a.

**Figure 4.**
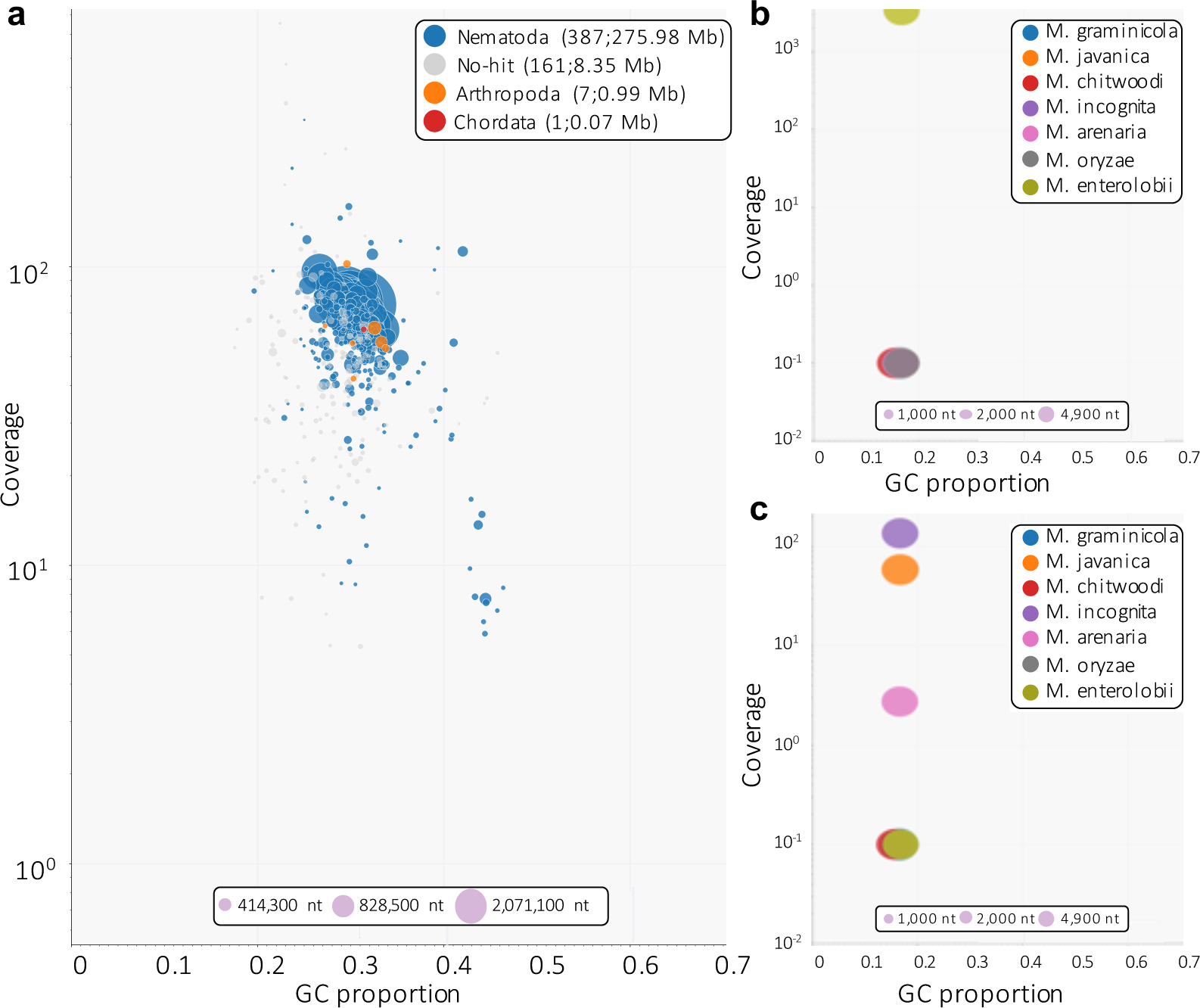
BlobPlot of different Meloidogyne genome assemblies. **A**. Blobplot showing taxonomic affiliation at the phylum rank level for the E1834 population of *M. enterolobii*. After removing contamination and mitochondrion, 556 contigs were left. The average GC content for *M. enterolobii* is equal to 30 ± 0.042. **B**. Mitochondrion only for the E1834 population, after removing contamination. No sign of other Nematoda within the assembly. **C**. Mitochondrion only for the previous *M. enterolobii* reference population provided by Koutsovoulos et al.^20^. No sign of *M. enterolobii* within the assembly. This BlobPlot revealed a contamination by other Meloidogyne spp..

Any contigs lacking taxonomic annotation are labeled as ‘no-hit’. For the non-Nematoda contigs falling perfectly inside the range of *M. enterolobii* GC content estimates^18^ (around 30%), a manual verification was conducted and eight of these contigs were kept. The proposed assignments from Blobtools were disregarded. Instead, for each of them we retained the highest-ranking result proposed by BLAST if the calculated percentage of identity surpassed 90%, the e-value did not exceed 1e-^50^, and the taxID belonged to the Nematoda phylum. This resulted in 556 final contigs for this assembly.

The Blobtools pipeline is a valuable tool for detecting possible contaminations in a genome assembly, especially those originating from distant species of different phyla. However, if the contamination comes from a closely related species with a comparable GC content or has been sequenced at a similar coverage, the classical approach will not detect a contamination. For this reason, we made slight adjustments to the methodology (Methods) to achieve a taxonomic classification based on different species within the Nematoda phylum, instead of between phyla only (Fig. 4b, 4c). We focused our analysis on different species within the Meloidogyne genus because (i) they are difficult to differentiate based on the morphology, (ii) they live in the same environment, (iii) they have similar GC content. Therefore, a non-negligible possibility for undetected contamination exists.

Using this modified BlobTools methodology, on the *M. enterolobii* population E1834 we have sequenced, we observed that the *M. enterolobii* reference mitochondrial genome from the NCBI was highly covered whereas all the other mitochondrial genomes from the other Meloidogyne species were not covered by our long reads. Hence, no evidence for contamination by other Meloidogyne species was found in the E1834 population (Fig. 4b).

For comparison, this method was applied to the previous long-read genome of *M. enterolobii*^20^, and surprisingly, it was found to be heavily contaminated by another *Meloidogyne* (Fig. 4c). Specifically, the *M. enterolobii* mitochondrial genome was not covered by the previous long reads while those of *M. incognita*, *M. javanica* and *M. arenaria* were all substantially covered. Approximately 60%, 30%, and 10% of the mitochondrial reads aligned with these mitochondrial genomes, respectively. Although this adjusted Blobtools approach suggested contamination of the previous Mma-II Swiss population from other root-knot nematodes, this alone was not sufficient to discriminate between these three closely related species.

Consequently, we combined this approach with SCAR markers. All the pairs of primers for the SCAR marker of the four Meloidogyne species of interest were aligned to the previous and current assemblies of *M. enterolobii*. Both for the L30 population of Burkina Faso and the E1834 population from Puerto Rico sequenced here, the pair of primers for the *M. enterolobii* SCAR marker matched the genome assemblies with 100% identity in the correct orientation on one single contig. This allowed identification of a virtual amplified sequence of 537bp, which is consistent with the ∼520bp estimated PCR product on the electrophoresis gel in Tigano et al^28^. In contrast, neither the *M. enterolobii* SCAR primers nor the reconstructed corresponding PCR product matched the previous Mma-II genome assembly, confirming the genome was probably not *M. enterolobii*. To further determine the possible source of contamination, we aligned the pairs of primers of the *M. incognita*, *M. javanica* and *M. arenaria* SCAR markers on the Mma-II genome. The *M. incognita* pair of primers matched perfectly on this previous assembly in the correct orientation and allowed reconstructing a virtual PCR product of 1192 bp, consistent with the estimated size of the PCR product of ∼1,200 bp for *M. incognita*^60^. Neither the pair of *M. incognita* primers nor the reconstructed PCR product matched the L30 or E1834 genome assemblies, and none of the *M. javanica* or *M. arenaria* pairs of primers matched any of the previously published or current *M. enterolobii* genomes.

Therefore, we can conclude that although no trace of contamination by closely related Meloidogyne species could be identified in the L30 or E1834 genome, there is clear evidence that the Mma-II population had been contaminated and replaced by *M. incognita*. The combination of SCAR marker analysis and a modification of Blobtools, specifically for ‘mitochondrion’, has resulted in a powerful tool for the examination and the verification of species purity.

### Genome completeness assessment

To evaluate the completeness of our genome assembly in terms of expected gene content among related species, we benchmarked nearly universal single-copy orthologs (BUSCO^68^ v5.2.2) by using the eukaryote_odb10 lineage dataset in fast mode. Despite the presence of a nematode dataset in BUSCO, it only contains seven species and none of them belong to the same clade as the root-knot nematodes. Therefore, we decided to use the more comprehensive Eukaryotic dataset, which encompasses 70 species. This procedure generates a report that indicates the number of genes that are universally or mostly conserved within the assembly and classifies them into several groups: complete, fragmented, single-copy, or duplicated. The results show that 71.4% (182/255) of BUSCO genes are complete and 12.5% are fragmented. This is a substantial improvement compared to the previously available assemblies (Table 5). Indeed, the Burkina Faso isolate of *M. enterolobii* reached eukaryotic BUSCO completeness score of 59.2% while the Mma-II assembly contaminated by *M. incognita* reached 69.4%.

**Table 5.** Ortholog BUSCO completeness analysis for different *M. enterolobii* using lineage dataset eukaryota_odb10. *Population contaminated by *M. incognita.* **This work.

| BUSCO<br>Categories | *M. enterolobii*<br>(L30<sup>69</sup>) | *M. enterolobii* *<br>(Swiss<sup>20</sup>) | *M. enterolobii*<br>(E1834) ** |
| --- | --- | --- | --- |
| Complete | 59.2% (151) | 69.4% (177) | 71.4% (182) |
| Single-copy | 29.8% (76) | 12.5% (32) | 14.9% (38) |
| Duplicated | 29.4% (75) | 56.9% (145) | 56.5% (144) |
| Fragmented | 18.4% (47) | 13.3% (34) | 12.5% (32) |
| Missing | 22.4% (57) | 17.3% (44) | 16.1% (41) |

BUSCO is a valuable and robust tool for assessing completeness in a genome assembly in terms of a widely conserved gene set. Nevertheless, in the case of less studied species, the analysis may lack precision if the newly assembled genome comprises variations not included in the initial BUSCO gene set, such as true copy number or sequence variants^40^. We then used Merqury once more to identify any copy-number errors and measure completeness and base accuracy via k-mers. Consequently, Merqury determined the proportion of reliable k-mers in the sequencing sample that were detected in the assembly, resulting in a completeness score of 99.60%. To establish Merqury’s base accuracy score, a binomial model for k-mer survival was employed, resulting in a Qscore of 65.70. Higher Qscores indicate a more precise consensus. For instance, Q30 corresponds to an accuracy of 99.9%, Q40 to 99.99%, and so on. In contrast, Burkina Faso and Swiss isolates have Qscores of 55.12 and 29.93, based respectively on their own reads.

### Gene prediction

Using the automated Eugene-EP pipeline, a total of 49,870 genes were predicted, with 45,924 being protein-coding genes and 3,946 being non-protein-coding genes such as rRNA, tRNA, and splice leader genes. These genes cover 84 Mb (approximately 29.48%) of the genome assembly length, with the exons spanning 44.51 Mb (around 15.60 %). On average, 5.26 exons are predicted per gene, and the gene length varies from a minimum of 150 bp to a maximum of 35,976 bp. The mean GC content is higher in either the protein-coding region (35.19%) or in the non-protein-coding gene regions (44.19%) compared to that of the whole genome (30.34%).

### Confirmation of genome structure and ploidy level

Although the *M. enterolobii* population E1834 genome has been predicted as a putative triploid based on k-mer analyses and Smudgeplot, it is important to further confirm the ploidy level of the genome assembly after annotation. The use of MCScanX reveals that a majority of gene duplicates create whole duplicated blocks, rather than dispersed independent duplications. Following the classification established by the duplicate_gene_classifier program implemented in the MCScanX package, 39,532 of the protein-coding genes (around 86.10%) are predicted to be duplicated at least once. As evidenced by Table 6, a majority of these coding genes (75.6%) show a duplication depth of two (meaning for these genes, two other copies exist), further reinforcing the idea that the genome is triploid. Furthermore, it was found that 69.76% of the protein-coding genes fall under the whole-genome duplication category of MCScanX, forming 516 syntenic blocks of collinear genes (see Fig. 5 for visualization of multiple syntenic blocks between different contigs). In addition, 12.61% of the genes are classified as dispersed duplicates, while 2.18% and 1.53% constitute proximal and tandem duplicates, respectively. These findings strongly suggest that the genome of *M. enterolobii* is triploid, confirming SmudgePlot results.

**Figure 5.**
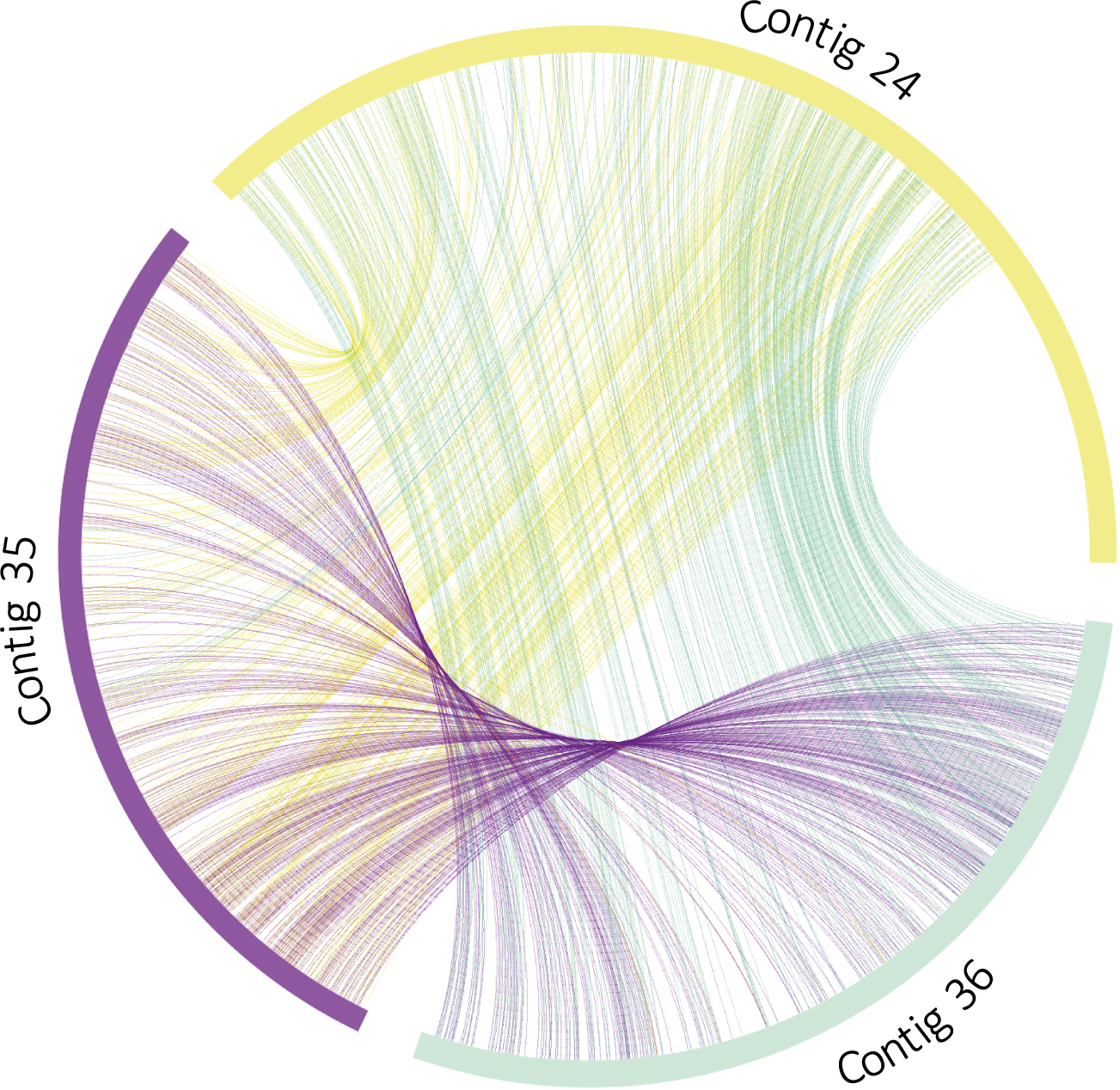
*M. enterolobii* exhibits a triploid genome. The circle plot produced by MCScanX shows collinear gene pairs forming homologous duplicated regions between three contigs. All the collinear gene pairs are linked with different curved colored lines between and within each contig.

**Table 6.** Duplicate gene classifier program of MCScanX for a self-comparison of *M. enterolobii*. Genes with a duplication depth of 0 are not duplicated, while a depth of 1 indicates a maximum of one copy, a depth of 2 indicates two copies, and so forth.

| Duplication depth | 0 | 1 | 2 | 3 | 4 | 5+ |
| --- | --- | --- | --- | --- | --- | --- |
| Gene numbers | 1992 | 4917 | 34720 | 2700 | 769 | 826 |
| Percentage (%) | 4.34 | 10.71 | 75.60 | 5.88 | 1.67 | 1.80 |

### Mitochondrial genome assembly and annotation

Using the Aladin package^44^, the mitochondrial genome of the *Meloidogyne enterolobii* population E1834 has been assembled and spanned a length of 19,193 bp with a GC content of 17.2% (Fig. 6). We have retrieved and annotated all the mitochondrially encoded subunits involved in the Mitochondrial respiratory chain, including the seven core subunits of the complex I, the cytochrome b of the complex III, the three cytochrome c, and the ATP synthase. We also obtained a full set of tRNAs, among which were in multiple copies (Ala, Ser, Leu and Asn) as well as ribosomal RNAs (rrnS, rrn12 and rrn16).

**Figure 6.**
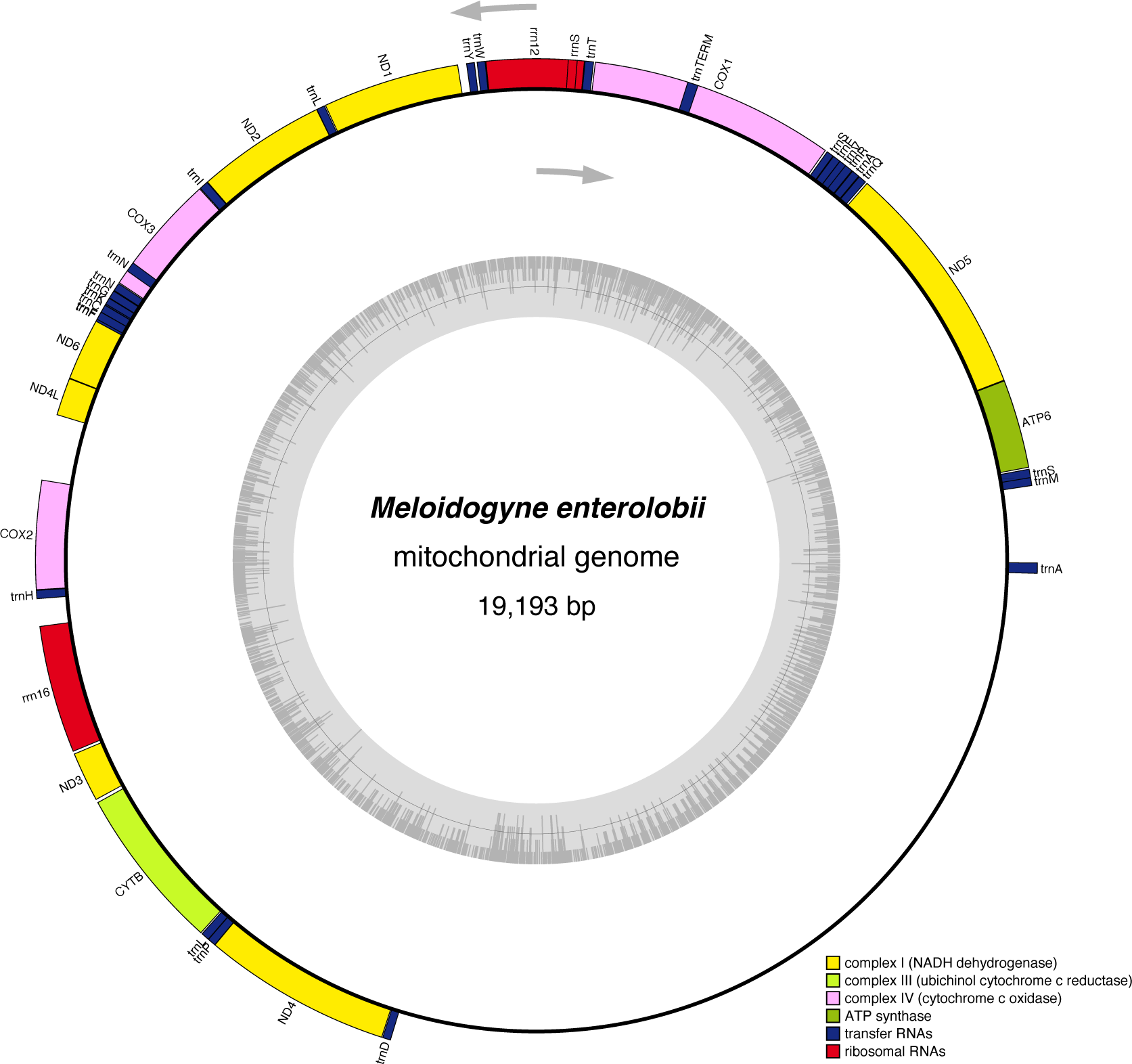
Mitochondrial genome organization of *M. enterolobii*. The inner circle displays the GC content while grey arrows denote the transcription direction. The rRNAs and tRNAs are respectively colored in red and blue. The various complexes of the CRM are represented in yellow, light green, pink, and dark green.

When blasted against the NCBI nt database, the reconstructed mitochondrial genome of the *Meloidogyne enterolobii* population E1834 returned as first hit the complete mitochondrial reference genome of *M. enterolobii*^45^, with 99.526% identity and an alignment length of 13,067 bp, as the primary hsp. The second-best hit corresponds to an incomplete mitochondrial genome from an *M. enterolobii* isolate discovered on sweet potatoes in the state of Carolina in the USA (GenBank: MW246173.1).

In contrast, reconstruction of the mitochondrial genome using Aladin on the Swiss population Mma-II yielded a ∼23kb genome which returned as first hit the *M. incognita* reference mitochondrial genome^58^ with >99% identity covering >97% of the query while the *M. enterolobii* reference mitochondrial genome only emerged as the fifth hit with only 87% identity covering 78% of the length.

These results further confirm the E1834 population we have sequenced is indeed *M. enterolobii*.

## Acknowledgements

We are grateful to the colleagues from the Netherlands Institute for Vectors, Invasive Plants and Plant Health (NIVIP) for providing the *M. enterolobii* population E1834 for this study. We thank the genotoul bioinformatics platform Toulouse Occitanie (Bioinfo Genotoul, https://doi.org/10.15454/1.5572369328961167E12) for providing computing resources. We are grateful to the bioinformatics and genomics platform, BIG, Sophia Antipolis (ISC plantBIOs, https://doi.org/10.15454/qyey-ar89) for computing and storage resources. We thank Claire Caravel for help in providing the primer sequences for SCAR identification of *Meloidogyne* species. Our *M. enterolobii* genome research is financially supported by a Franco-German bilateral grant ANR-DFG “AEGONE”, reference ANR-19-CE35-0017 and reference No 431627824.

## Author contributions

E.G.J.D. and S.K. conceived the research idea and acquired the funding. E.G.J.D. supervised all the bioinformatics analyzes, performed SCAR marker virtual PCRs, contributed to manuscript writing and reviewing. S.K. supervised all the nematode rearing and DNA extraction experiments, contributed to manuscript writing and reviewing. M.P. performed reads processing, genome assembly, contamination and purity check, completeness assessment, ploidy and genome size and structure estimation and wrote the manuscript. H.G. generated single egg mass lines, performed maintenance of the nematode collection, DNA extraction experiments, and contributed to manuscript writing. C.R. performed gene prediction and wrote the corresponding method section. M.S., C.L.R., and J.L. performed library and PACBIO HiFi sequencing and contributed to manuscript writing.

